# LIN-35 is necessary in both the soma and germline for preserving fertility in *Caenorhabditis elegans* under moderate temperature stress

**DOI:** 10.1101/2022.02.28.482413

**Authors:** Brian P. Mikeworth, Frances V. Compere, Kristen Quaglia, Lisa N. Petrella

**Author notes:** Department of Biology, Syracuse University, Syracuse, New York, United States of America (Frances V. Compere). Corresponding author, (LNP).

## Abstract

Maintenance of germline function under stress conditions is crucial for species survival. The germline across many species is especially sensitive to elevated temperature stress. We have investigated the role that the pocket protein LIN-35 plays in preserving fertility in *C. elegans* under moderate high temperature stress. Here we show that *lin-35* mutants have a number of temperature sensitive germline defects. The decreased brood size seen at elevated temperatures is more severe in *lin-35* mutants than wild type. This loss of fertility under temperature stress is primarily due to loss of zygotic but not maternal LIN-35. Additionally, we have found that expression of LIN-35 is necessary in both the germline and soma for the preserving fertility at elevated temperature. Specifically, germline expressed LIN-35 is primarily what is important for maintaining fertility in hermaphrodites, but broad somatic expression of LIN-35 is also necessary for oocyte formation and/or function at elevated temperatures. Additionally, we found that overexpression of LIN-35 in the germline only can stabilize the formation of P granules in primordial germ cells. Our data adds to the emerging critical role that LIN-35 plays in buffering tissues against stress.

## INTRODUCTION

LIN-35 is the single *C. elegans* pocket protein with homology to the three mammalian pocket proteins including Rb (retinoblastoma), p107, and p130 [1]. Pocket proteins generally participate in transcriptional repression, which in mammals has been linked both to regulation of the cell cycle and cell fate decisions [2–3]. In *C. elegans*, LIN-35 is expressed in almost all tissues and has distinct roles in many of these tissues including repression of germline expressed genes in the soma, suppression of nuclear divisions in the intestine, repression of RNAi pathways in the soma, and regulation of apoptosis in the germline [5–7]. *lin-35* mutants also demonstrate a range of synthetic phenotypes, which only occur in the absence of another factor. These synthetic phenotypes demonstrate that LIN-35 also plays some role in regulating vulval cell fate specification, somatic apoptosis, and somatic gonad development [8–10]. Despite its role in so many systems, *lin-35* mutants are viable and fertile at moderate temperatures, although they demonstrate slow growth and reduced brood sizes [5, 11]. Loss of LIN-35 leads to substantial changes in gene expression in the germline, suggesting that the loss of fertility in *lin-35* mutants is at least in part intrinsic to germ cells [12]. However, loss of LIN-35 also has effects on the somatic gonad and intestine [4–5, 8, 13], both of which contribute to fertility [14–15]. Therefore, it is not yet clear if loss of LIN-35 in these supporting somatic tissues also contribute to the reduced brood size of *lin-35* mutants.

Fertility is temperature sensitive across many animal species, including in *C. elegans*. Wildtype fertility peaks around 18-20°C and shows a decline as temperatures increase to 26°C [16–17]. Between 26°C and 27°C *C. elegans* worms become essentially sterile [18–19]. As temperature increases, the loss of fertility appears to start with a decrease in the number of functional sperm followed by a loss of oocyte function at slightly higher temperatures [17–19]; however, the specific mechanisms that are failing at different temperatures are poorly understood. Interestingly, many *lin-35* mutant phenotypes are also temperature sensitive, including the synthetic multivulva phenotype, larval growth (high temperature larval arrest phenotype), misregulation of developmental chromatin compaction, and ectopic germline gene expression [13, 20–21]. Previous work has shown that the fertility of *lin-35* mutants is decreased at 25°C compared to 20°C [5]. Since wildtype fertility is also decreased at this temperature, it is unclear if this represents a temperature sensitive phenotype or just the same trend in *lin-35* mutants as is seen in wild type.

Here we show that *lin-35* mutants have a temperature sensitive fertility defect. Using somatically expressed LIN-35(+) transgenes to bypass the high temperature larval arrest phenotype, were discovered to that *lin-35* mutants have a temperature sensitive loss of fertility that is greater than the loss of fertility in wildtype animals. At elevated temperatures, *lin-35* mutants also have defects in oocyte formation. Finally, we show that while the germline is the primary tissue where *lin-35* expression is necessary for fertility, *lin-35* expression in somatic tissues also plays a role in maintaining fertility.

## METHODS

### Strains and Experimental Design

*C. elegans* were cultured on NGM plates seeded with the *Escherichia coli* strain AMA1004 at 20°C unless otherwise noted. The following strains were used: N2 (Bristol) was used as wild type, MT10430 *lin-35(n745)* (CGC), LNP0044 *petEx1*[let-858p::lin-35::GFP + rol-6], LNP0043 *lin-35(n745)*; *petEx1[let-858p::lin-35::GPF + rol-6]*, SS0991 *bnEx56[elt-2p::lin-35::GFP + rol-6]* (Petrella *et al*., 2011), LNP0016 *lin-35(n745); bnEx56[elt-2p::lin-35::GFP + rol-6*], LPN0031 *vrIs56 [pie-1p::lin-35::GFP::FLAG::lin-35 3’UTR + unc-119(+)]* (transgene from Kudron et al., 2013 obtained from the CGC), LNP0022 *lin-35(n745); vrIs56 [pie-1p::lin-35::GFP::FLAG::lin-35 3’UTR + unc-119(+)]* LNP0038 *vrIs93[mex-5p::lin-35::GFP::FLAG::lin-35 3’UTR + unc-119(+)]* (transgene from Kudron et al., 2013 obtained from the CGC), LNP0023 *lin-35(n745); vrIs93[mex-5p::lin-35::GFP::FLAG::lin-35 3’UTR + unc-119(+)]*, LNP0042 *+/hT2 [bli-4(e937) let-?(q782) qIs48] (I;III)* SS0911 *lin-35(n745) /hT2 [bli-4(e937) let-?(q782) qIs48] (I;III)* (Petrella et al., 2011), and LW697 *ccIs4810 [lmn-1p::lmn-1::GFP::lmn-1 3’utr + (pMH86) dpy-20(+)]* (CGC). Some strains (as noted) were provided by the CGC, which is funded by NIH Office of Research Infrastructure Programs (P40 OD010440).

#### Transgenes

We generated a strain containing pan-somatic expression of LIN-35(+) tagged with GFP from the same plasmid used to create the intestine-specific expression of LIN-35(+) by swapping the 5’ promoter sequences (*elt-2p* to *let-858p)* using the 2100 base pairs directly upstream of the *let-858* gene. Wildtype (N2) strains were microinjected with the *let-858p::lin-35::GFP* plasmid and pRF4 to create the extrachromosomal array *petEx1[let-858p::lin-35::GFP* + *rol-6]*. lin-*35(n745)* mutants were confirmed via random fragment length polymorphism (RFLP) and western blot analysis.

#### Temperature treatments

Three temperature treatments were used in this study. (1) Continuous exposure to 20°C: experiments were performed on strains maintained continuously at 20°C. (2) Continuous exposure to 26°C: P0 hermaphrodites were up-shifted to 26°C at the L4 stage and experiments were done on F1 hermaphrodites that had experienced their entire lifespan at 26°C (3) Up-shift from 20°C to 26°C: hermaphrodites were developed at 20°C until the L4 larval stage and then up-shifted to 26°C and experiments were done on the upshifted P0 worm.

#### Western Blot Analysis

Eighty F1 progeny raised at either 20°C or 26°C were collected for total protein samples in wild type, *lin-35(n745)*, and strains containing somatic *lin-35::GFP* transgenes. Total protein from the entire sample was run on a 7.5% SDS-PAGE gel and transferred to a Nitrocellulose membrane. Membranes were blocked (5% non-fat dry milk in PBS) and incubated in 1:1000 Rabbit anti-LIN-35 [22] in block and 1:200 Mouse anti-TUBB-E7 (Developmental Studies Hybridoma Bank, University of Iowa, Iowa City, IA, USA) in block overnight at 4°C. Membranes were blocked a second time and incubated with 1:2000 Goat anti-rabbit-HRP (ThermoFischer Scientific, USA) and 1:2000 Goat anti-mouse-HRP (ThermoFischer Scientific, USA) conjugated antibodies for one hour in the dark at room temperature. Blots were visualized via HRP-chemiluminesence (Amersham RPN2232) and exposure to X-ray film.

#### Brood Size Assay

To assay brood size, 10-15 L4 hermaphrodites were placed on plates individually and kept at their respective temperatures. Each hermaphrodite assayed was moved to fresh plates daily until no embryos were seen on plates. F2 progeny were counted during the L4 larval stage or as young adults. Brood size assays of mated hermaphrodites were conducted following a similar protocol except that 3 N2 males raised at 20°C were added to the plate. All hermaphrodites and males were transferred to seeded NGM plates daily, with the addition of three N2 males raised at 20°C on day 3 post-initial mating. Statistical analysis was performed with exclusion of sterile hermaphrodites using either two-way ANOVA or students T-test in GraphPad PRISM (GraphPad Software Inc., La Jolla, CA, USA).

#### Maternal Effect

Maternal effect of LIN-35 was observed through the use of the hT2 balancer tagged with GFP. P0 *lin-35(n745)/hT2::GFP* where plated at 20°C and upshifted to 26°C at the L4 larval stage. By cloning out *lin-35(n745)/hT2::GFP* or *lin-35(n745)* F1 progeny at the L4 larval stage, we were able to test M+Z+ and M+Z-respectively. To test for M-Z+, *lin-35(n745)* mothers were mated to LW697 *ccIs4810 [lmn-1p::lmn-1::GFP::lmn-1 3’utr + (pMH86) dpy-20(+)]* males at 26°C, where GFP(+) F1 M-Z+ progeny were selected and clone out at the L4 larval stage. Brood size for each cloned out F2 was counted as described above.

#### Immunocytochemistry

For L1 staining: P0 mothers were moved onto a new plate at the L4 stage and either maintained at 20°C or up-shifted to 26°C. After ~24 hours, young adult P0 mothers were placed in 1XM9 solution without bacteria and allowed to lay embryos overnight at either 20°C or 26°C. F1 L1 larvae were transferred to slides coated with poly-L-Lysine, freeze cracked, and fixed with methanol/acetone [23]. For adult germlines: P0 worms were moved onto a new plate at the L4 stage and either maintained at 20°C or upshifted to 26°C. F1 worms were moved onto a new plate at the L4 stage and maintained at previous temperature. After ~24 hours F1 adults were transferred without bacteria to slides coated with poly-L-Lysine in 0.1 mM levamisole. Germlines were dissected before being freeze cracked, and fixed with methanol/acetone. Samples were blocked (1.5% BSA, 1.5% OVA, 0.05% NaN_3_ in PBS) and incubated with primary antibody 1:10,000 rabbit anti-PGL1 [24] overnight at 4°C. Samples were incubated with secondary antibodies conjugated with Alexa Fluor 568 or 488 at 1:300 (ThermoFischer Scientific, USA) in PBS for 1 hour at room temperature. Slides were treated with 5mg/ml DAPI in 50 mL PBS for 10 min, washed 3 times with PBS for 10 min at room temperature and mounted on gelutol mounting medium. Z-stacks were taken using Nikon A1R Inverted Microscope Eclipse Ti confocal microscope with NIS Elements AR 3.22.09 at 60X. For L1s, images were blinded and then scored for PGL-1 localization.

#### Sperm Localization Assay

Sperm localization was observed by mating N2 males stained with MitoTracker Red CMXRos (Invitrogen) to mutant hermaphrodites. P0 mothers were plated at 20°C or upshifted to 26°C at the L4 larval stage. Males raised at 20°C were stained with 50μL of 50μM Mitotracker Red CMXRos in M9 for 2-4 hours in the dark. 25 stained males were placed onto NGM plates seeded with HB101 *E. coli* containing approximately 20 F1 L4 larvae (raised at 20°C or 26°C) and allowed to mate for 12-16 hours. Mated hermaphrodites were transferred to a clean plate for 1 hour before imaging to allow mated sperm to settle. Worms were imaged on a 2% agarose pad in 10mM Levamisole. DIC and fluorescent images were acquired using a Nikon Eclipse TE2000-S (Nikon Instruments Inc., Elgin, IL, USA) inverted microscope at 60X with Q-Capture Pro (QImaging, Surrey, BC, Canada) imaging software. Sperm were scored as either primarily localizing within the spermatheca, a mixture of localization to the uterus and spermatheca, or primarily localized within the uterus.

#### Oocyte Counting

Adult hermaphrodites were imaged live using Normarski optics DIC and images were acquired using a Nikon Eclipse TE2000-S (Nikon Instruments Inc., Elgin, IL, USA) inverted microscope at 60X with Q-Capture Pro (QImaging, Surrey, BC, Canada) imaging software. One gonad arm was imaged per hermaphrodite and scored for the presence of one or more than one oocyte in the proximal gonad. Oocytes were defined as cells having a large clear nucleus and taking up the entire width of the gonad.

## RESULTS

### Somatically expressed lin-35 can rescue high temperature arrest and allow for germline temperature sensitivity analysis

We wanted to know if germline defects in *lin-35* mutants may be temperature sensitive in a manner similar to somatic *lin-35* phenotypes [13, 20–21]. However, germline development and function occur primarily after the L1 stage, thus the ~100% high temperature larval arrest (HTA) seen in *lin-35* mutants at 26°C precluded the study of the effect of temperature on the adult germline in *lin-35* mutants directly. Therefore, we used two transgenic strains that rescue high temperature larval arrest, but still lack *lin-35(+)* expression in the germline (Fig 1a). First, we expressed *lin-35::GFP* using the *elt-2* promoter (*elt-2p::lin-35::GFP*), which allows for intestinal specific expression of *lin-35(+)*. The *elt-2p::lin-35::GFP* transgene in a *lin-35* mutant background rescued the HTA phenotype [13], while leaving all non-intestinal tissues mutant for *lin-35* function. We also created a transgenic line that expressed *lin-35::GFP* in all somatic cells under the ubiquitous *let-858* promoter (*let-858p::lin-35::GFP*). We found that expression of pan-somatic *let-858::lin-35::GFP* also rescued the HTA phenotype (data not shown). Both of these *lin-35(+)* transgenes are on extrachromosomal arrays, which generally are silenced in the germline independent of the promoter present [25]. Our analysis of both LIN-35(+) and GFP expression showed no detectible expression in the germline (data not shown). These two *lin-35(+)* transgenes allowed us to analyze the function of *lin-35* in the germline at high temperature.

**Figure 1:**
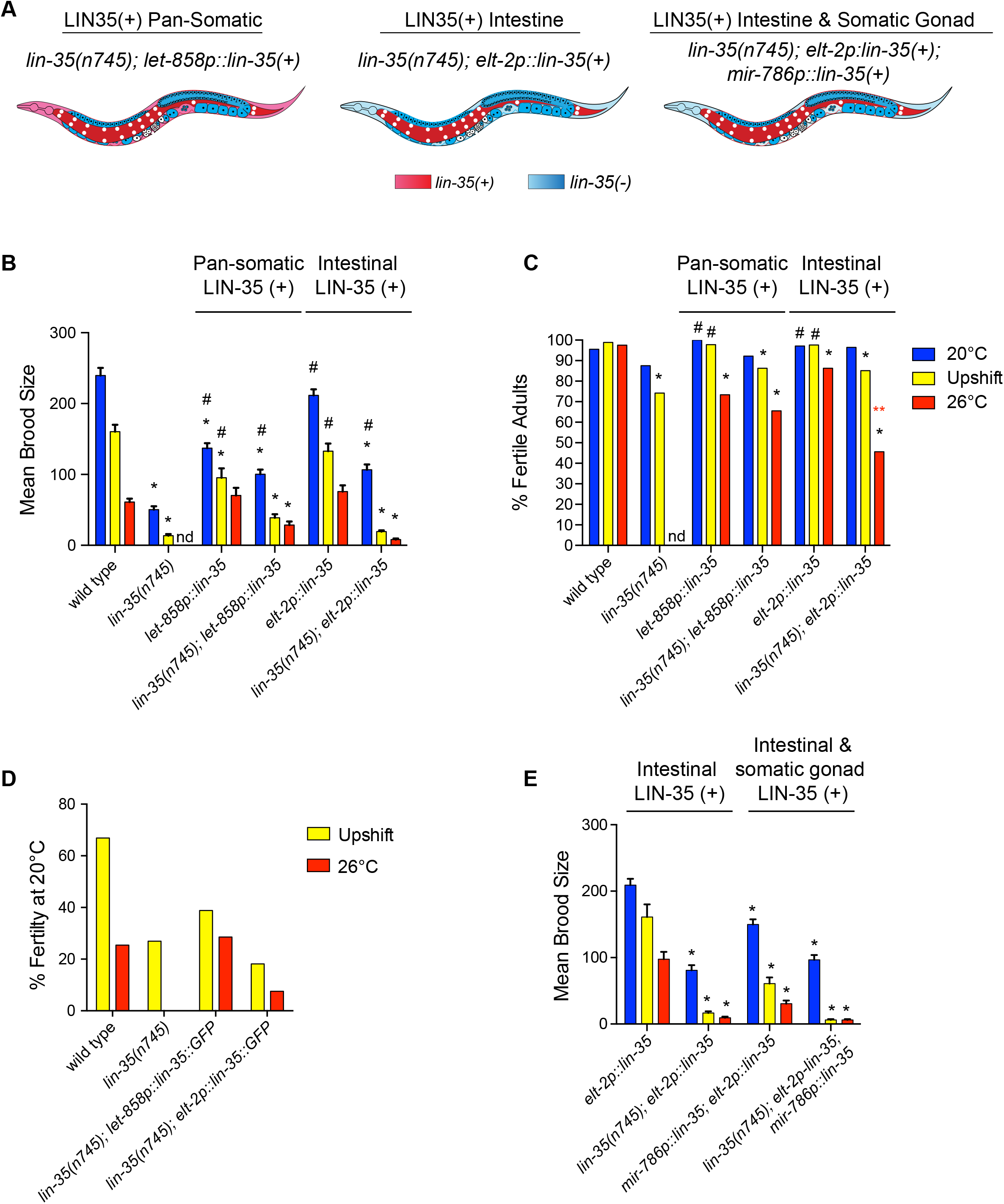
Somatic expression of *lin-35(+)* weakly rescues *lin-35* mutant fertility defects. (A) Schematic of different somatic transgenes used in this study. (B) Mean brood size of fertile worms for each treatment. Expression of *lin-35(+)* from either somatic transgene only rescues fertility at 20°C. nd: no data, * significantly different than wild type at the same temperature, # significantly different than *lin-35* mutants at the same temperature. *P*-value ≤ 0.05 using two-way ANOVA with Tukey correction. Error bars indicate ± SEM. (C) Percentage of hermaphrodites that are fertile for each treatment. *lin-35* mutants expressing intestinal *lin-35(+)* have significantly fewer fertile worms at 26°C than those expressing pan-somatic *lin-35(+)*. * significantly different than wild type at the same temperature, # significantly different than *lin-35* mutants at the same temperature, ** (red) significantly different than *lin-35(n745); let-858p::lin-35 P*-value ≤ 0.05 using a Fisher’s exact test. (D) The decrease in fertility represented by the percentage of the mean brood size with upshift or at 26°C compared to 20°C within the strain. With upshift, *lin-35* mutants retain less fertility than wild type even if expressing somatic *lin-35(+)*. At 26°C, *lin-35* mutants with pan-somatic but not intestinal expression of *lin-35(+)* retain a similar level of fertility as wild type. (E) Mean brood size of fertile worms for each treatment with expression of *lin-35(+)* in the somatic gonad. * significantly different than *elt-2p::lin-35* at the same temperature. *P*-value ≤ 0.05 using two-way ANOVA with Tukey correction. Error bars indicated ± SEM. Temperature treatments include continual growth at 20°C (blue), upshift from 20°C to 26°C at the L4 stage (yellow), and continual growth at 26°C (red).

### *lin-35* mutants have temperature sensitive loss of fertility that is not rescued by somatic expression of *lin-35::GFP*

To determine if *lin-35* mutants demonstrate temperature sensitive germline defects, we scored both the brood size of individual worms and the percentage of fertile hermaphrodites in the population with different temperature treatments when *lin-35* was missing from the germline. *lin-35* mutants have been previously shown to have a reduced brood size at 20°C; however, the cause of the reduction is unknown [5, 11–12]. We also saw that *lin-35(n745)* mutants with no *lin-35(+)* somatic transgene have a drastically reduced brood size; however, the percentage of fertile worms in the population is only weakly reduced compared to wild type (Fig 1b-c). *lin-35* mutants expressing either somatic *lin-35(+)* transgene at 20°C showed a partial but significant rescue of fertility defects, including an increase in both brood size and the number of fertile worms in the population (Fig 1b-c). These data indicate that part of the loss of fertility in *lin-35* mutants at 20°C is due to loss of *lin-35* function in somatic tissues.

As temperature increases to 26°C brood size decreases in wildtype *C. elegans* hermaphrodites, but the percentage of hermaphrodites that are fertile remains close to 100% [18–19]. To allow for comparison of transgenic lines with *lin-35(n745)* mutants with no *lin-35(+)* transgene, we grew hermaphrodites at 20°C until the L4 stage when they were upshifted to 26°C (Upshift). Because the exposure to 26°C occurred after most of larval development, this allowed us to bypass the HTA phenotype seen in *lin-35* mutants. Similar to what is seen with either type of temperature stress in wild type worms, we saw a significant decreased brood size in *lin-35* mutants with or without a somatically expressed *lin-35(+)* transgene, while the majority of the hermaphrodites maintained fertility (Fig 1b-c). Because the different genotypes had different brood sizes at 20°C, we compared relative brood size of each genotype upon upshift to that same strain’s brood size at 20°C (Fig 1d). We found that *lin-35* mutants with or without somatic *lin-35(+)* transgenes had a stronger relative loss of fertility when upshifted to 26°C than wild type (Fig 1d). Overall, the fertility of *lin-35* mutants was more sensitive to increases in temperature at the L4 stage than wild type, and expression of wildtype *lin-35* in the soma did not rescue this temperature sensitivity.

We next investigated the effects on fertility of raising hermaphrodites for their entire development at 26°C. While *lin-35* mutants expressing either somatic *lin-35(+)* transgene showed similar effects on brood size when animals were raised at 20°C, whether upshifted or not, there was a marked difference in brood size between the *lin-35(+)* transgenes in *lin-35(n745)* mutants raised at 26°C. At 26°C, significantly fewer *lin-35* mutants that expressed the intestinal *lin-35(+)* transgene were fertile than those that expressed the pan-somatic *lin-35(+)* transgene (Fig 1c). Additionally, the relative decrease in brood size between worms raised at 20°C and 26°C was greater in *lin-35* mutants that expressed the intestinal *lin-35(+)* transgene than those that expressed the pan-somatic *lin-35(+)* transgene (Fig 1d). In fact, *lin-35* mutants that expressed pan-somatic *lin-35(+)* showed a reduction in brood size indistinguishable from wild type.

The partial rescue of fertility at 26°C in *lin-35* mutants with the pan-somatic *lin-35(+)* transgene but not the intestinal *lin-35(+)* transgene suggested that expression of *lin-35(+)* in non-intestinal somatic lineages is important when worms are raised at 26°C. *lin-35* has been previously shown to function in the somatic gonad [8]. Therefore, we expressed *lin-35::GFP* under the *mir-786* promoter, which expresses in the somatic gonad [26], in combination with the intestinal expression of *lin-35(+)* transgene. However, there was no increase in brood size at elevated temperature when there is expression of *lin-35(+)* in the somatic gonad (Fig 1e). As even pan-somatic expression of *lin-35(+)* resulted in a much weaker rescue at 26°C than at 20°C, there is likely a germline intrinsic function for LIN-35 that is necessary for preserving fertility at elevated temperatures.

In the course of these experiments, we observed that pan-somatic expression of *lin-35(+)* in a wildtype background (i.e. somatic overexpression of *lin-35(+))* resulted in a reduced brood size compared to wild type without the *lin-35(+)* transgene (Fig 1b). In a *lin-35* mutant background, this *lin-35(+)* transgene fully rescues the HTA phenotype and rescues fertility close to the level of the *lin-35(+)* transgene in the wildtype background. One possible explanation for the reduced brood size of when *lin-35(+)* is expressed in wildtype background is that the level of overexpression of LIN-35::GFP driven by the strong *let-858* promoter may have a partially dominant negative affect in a somatic tissue that interacts with the germline. Western blots showed that in wildtype animals expressing *let-858p::lin-35(+)::GFP* the level of native LIN-35 was reduced (S1 Fig). Previous work has shown that LIN-35 can bind its own promoter [22, 27], thus it is possible that the expression of LIN-35::GFP may down-regulated the native LIN-35 locus.

### Germline rescue of *lin-35:GFP* rescues temperature effects on fertility

To determine if there is germline intrinsic function for LIN-35 in preserving fertility at 26°C, we investigated the temperature effects on fertility of germline-specific expression of *lin-35* using two different germline promoters: *pie-1p::lin-35::GFP* and *mex-5p::lin-35::GFP* [28]. Because somatic expression of *lin-35* is necessary to rescue the HTA phenotype, we could only examine the effects of temperature in upshifted L4 hermaphrodites. Expression of *lin-35(+)* from either transgene resulted in a strong partial rescue of brood size at both 20°C and upon upshift from 20°C to 26°C (Fig 2a). Additionally, the relative decrease in brood size between worms upshifted to 26°C and worms raised at 20°C for either germline-specific *lin-35(+)* transgene was equivalent to wild type (Fig 2b). This is in contrast to soma-only expression of *lin-35(+)* that was able to partially rescue absolute brood size but still had a greater relative decrease in brood size when challenged with elevated temperature (Fig 1d). Therefore, germline intrinsic expression of LIN-35 is primarily responsible for the preserving temperature effects on brood size.

**Figure 2:**
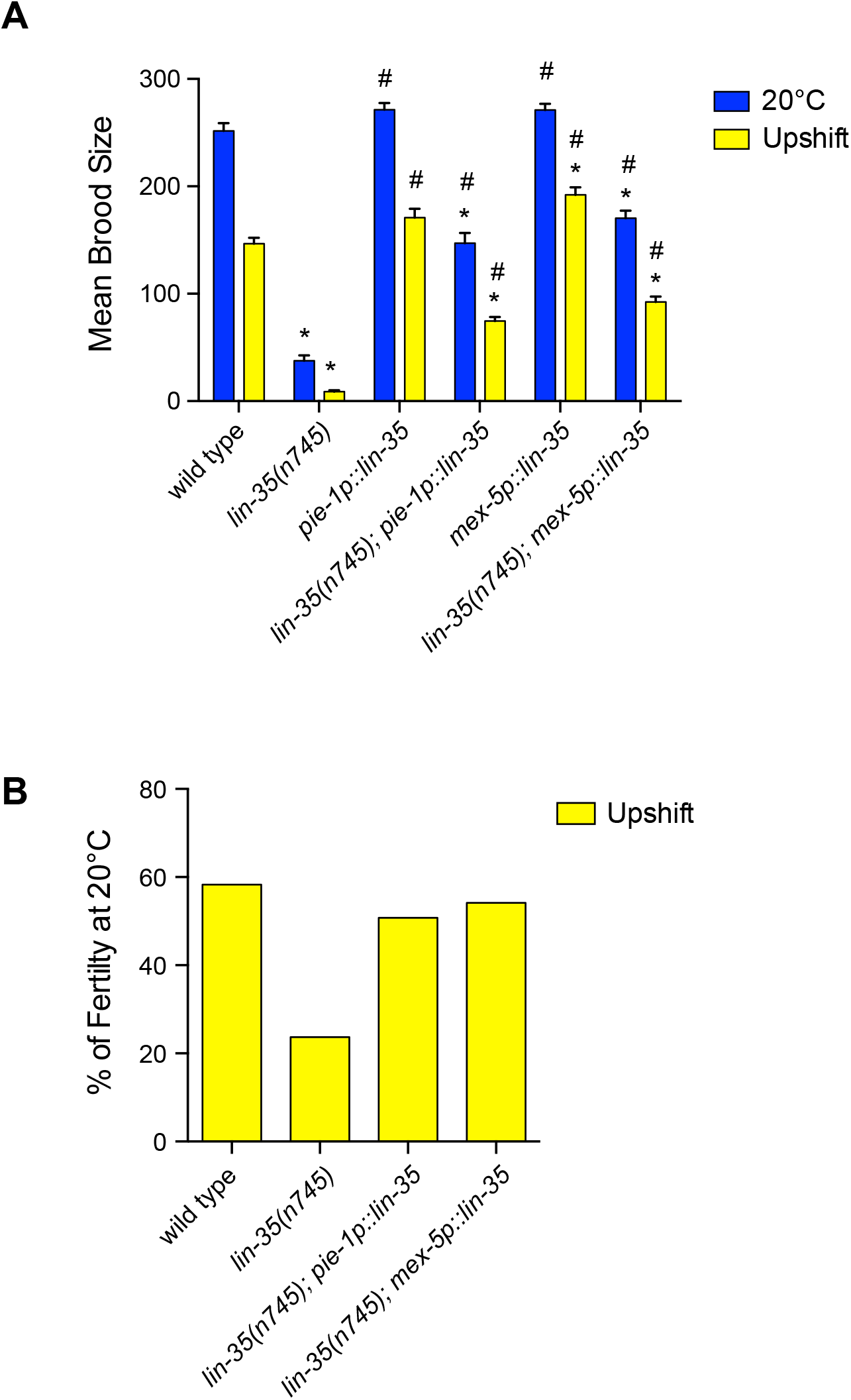
Germline expression of *lin-35(+)* strongly rescues *lin-35* mutant fertility defects. (A) Mean brood size of fertile worms for each treatment. Expression of either germline *lin-35(+)* transgene significantly increases brood size at both 20°C and with upshift. * significantly different than wild type at the same temperature, # significantly different than *lin-35* mutants at the same temperature. *P*-value ≤ 0.05 using ANOVA with Turkey correction. Error bars indicated ± SEM. (B) The decrease in fertility represented by the percentage of the mean brood size with upshift compared to 20°C within the strain. With upshift *lin-35* mutants expressing germline *lin-35(+)* retain a similar level of fertility as wild type. Temperature treatments include continual growth at 20°C (blue) and upshift from 20°C to 26°C at the L4 stage (yellow).

### Loss of zygotic *lin-35* is sufficient to cause sterility at high temperature

We wanted to know if the increased loss of fertility at 26°C was a maternal effect phenotype like the HTA phenotype [13] and the synMuv phenotype [9]. We tested the fertility of first generation *lin-35(n745)* homozygous mutants from heterozygous mothers where *lin-35(n745)* is balanced by the *hT2* translocation that expresses GFP in the pharynx (M+Z-). The hT2 translocation itself causes a high penetrance of embryonic lethality in a wildtype background [27]. Therefore, we compared *lin-35(n745)* M+Z-hermaphrodites to wildtype worms that were the first generation GFP-progeny of +/hT2 hermaphrodites (Fig 3A). At 20°C *lin-35(n745)* M+Z-homozygotes that only have maternally loaded *lin-35(+)* have an intermediate brood size between wild type and *lin-35(n745)* M-Z-homozygous mutant hermaphrodites (Fig 3A). Thus, *lin-35* mutants at 20°C demonstrate partial maternal effect fertility defects. However, at 26°C *lin-35(n745)* M+Z-hermaphrodites were completely sterile. Thus, the loss of solely zygotic *lin-35(+)* expression is sufficient to result in a total loss of progeny production at 26°C.

**Figure 3:**
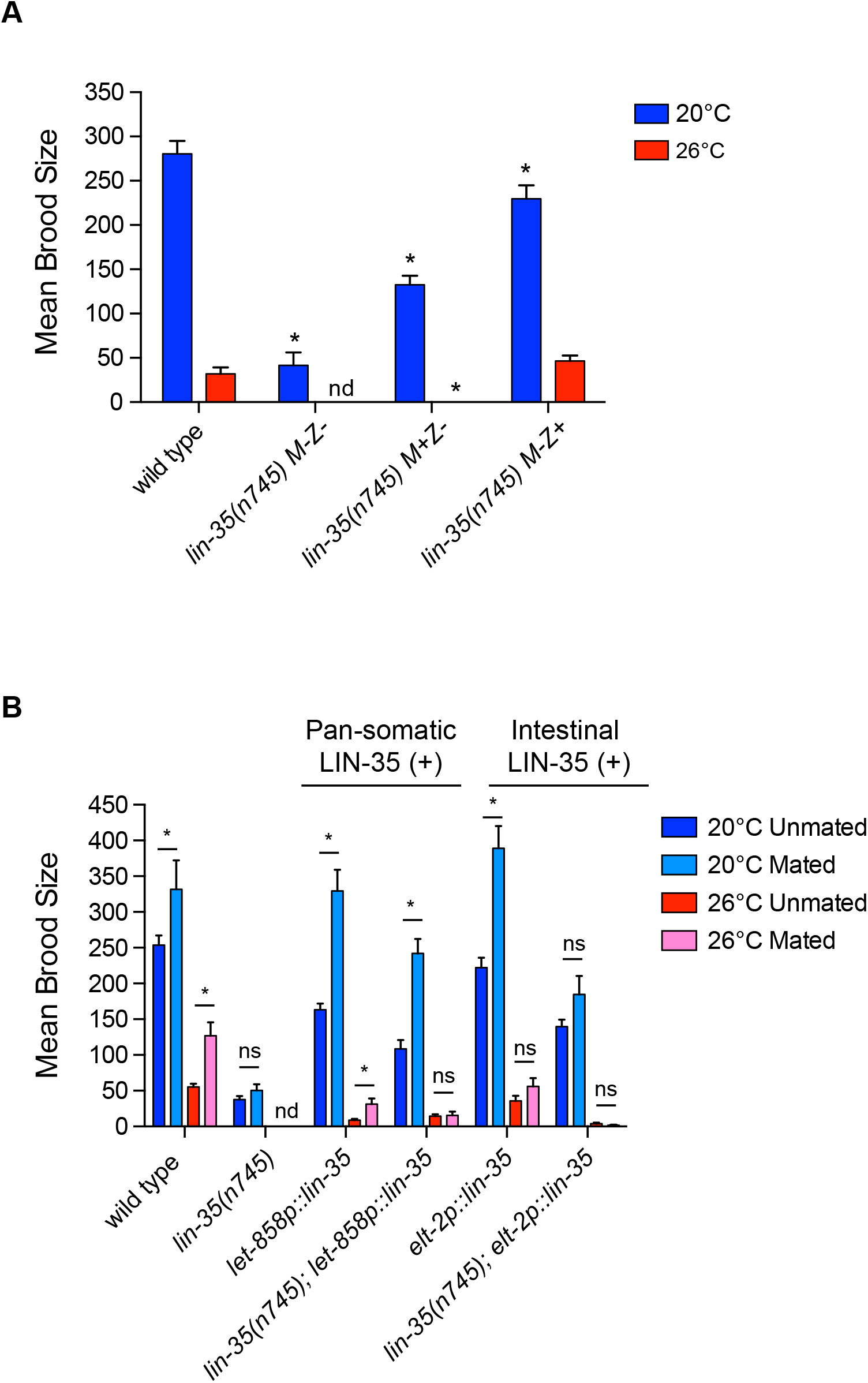
Fertility defects in *lin-35* mutants are not maternal effect and are not strongly rescued by mating at high temperature. (A) Mean brood size of fertile worms for each treatment. Maternal load of *lin-35(+)* is not sufficient for fertility to be maintained in *lin-35* M+Z-mutants at 26°C and zygotic expression of *lin-35* in M-Z+ mutants rescues fertility loss in mutants at 26°C. M-Z-animals do not inherit any wildtype *lin-35* products (protein or RNA) from their mother and do not have a wildtype copy of the *lin-35* gene, M+Z-animals do inherit *lin-35* products from their mother but do not have a wildtype copy of the *lin-35* gene, and M-Z+ do not inherit any wildtype *lin-35* products (protein or RNA) from their mother but do have a wild type copy of the *lin-35* gene. * significantly different than wild type at the same temperature. *P*-value ≤ 0.05 using two ANOVA with Turkey correction. Error bars indicated ± SEM. (B) Mean brood size of fertile worms for each treatment. Mating with wildtype males does not significantly increase the brood size of *lin-35* mutants expressing either somatic transgene at high temperature. * significantly different from same strain unmated at the same temperature. *P*-value ≤ 0.05 using students t-test. Error bars indicated ± SEM. Temperature treatments include continual growth of hermaphrodites at 20°C (blue) and continual growth at 26°C (red). For mated experiments, males were always grown to adulthood at 20°C and then mating occurred at the temperature of the hermaphrodite growth.

To further investigate the role of zygotically expressed *lin-35* in fertility we crossed *lin-35(n745)* M-Z-hermaphrodites with wildtype males that carried a GFP marker and scored the brood size of resulting *lin-35* heterozygote M-Z+ hermaphrodites. These worms lacked maternally loaded LIN-35(+) protein and mRNA needed in early embryogenesis, but expressed zygotic LIN-35(+) protein. At 20°C *lin-35(n745)* M-Z+ hermaphrodites had a significantly smaller average brood size than wildtype hermaphrodites, but a significantly higher average brood size than *lin-35(n745)* M+Z-hermaphrodites (Fig 3A). At 26°C *lin-35(n745)* M-Z+ hermaphrodites have an average brood that was not significantly different than wildtype hermaphrodites (Fig 3A). Thus, zygotic expression alone can strongly rescue fertility of *lin-35* mutants at low temperature and completely rescue fertility at high temperatures.

### Mating does not significantly rescue fertility in *lin-35* mutants under most conditions

Decreased brood size in wildtype worms at 26°C is primarily due to decreased sperm function [17–19]. To determine if the significant loss of brood size in *lin-35* mutants at 26°C is also primarily due to decreased sperm function, we crossed *lin-35(n745)* hermaphrodites with and without *lin-35(+)* somatic transgenes to wildtype males raised at 20°C to provide functional sperm. At 20°C, the number of sperm is the limiting factor in brood size [30]; therefore, crossing with a male which provides more sperm generally leads to an increase in brood size. For hermaphrodites grown at 20°C there was a significant increase in brood size in all wildtype strains when mated; however, only *lin-35(n745)* mutants expressing the pan-somatic *lin-35(+)* transgene showed an increase in brood size when mated (Fig 3b). At 26°C the number of functional sperm made by the hermaphrodite has been shown to be decreased; therefore, crossing with a male raised at 20°C should also increase brood size if oocyte function is not compromised. For hermaphrodites grown at 26°C, neither strain expressing a somatic *lin-35(+)* transgene in *lin-35(n745)* mutants showed an increase in brood size (Fig 3b).

There are two possible scenarios that could lead to the observation that *lin-35* mutants with or without somatic *lin-35(+)* transgene expression show limited increases in brood size when mated. First, when *lin-35* mutant hermaphrodites mate with males there could be limited sperm transfer/migration to the spermatheca. Second, *lin-35* mutants could have a defect in oocytes that limits the ability of male sperm to create cross progeny [31]. To test the first scenario, we labeled male sperm by incubating males with mitoTracker Red and then mated them with hermaphrodites for a period of 12-16 hours. We then assessed if sperm were present in hermaphrodites and able to migrate to the spermatheca (Fig 4a-b). We found in most strains for hermaphrodites raised at either 20°C or 26°C there was strong localization of male sperm to the spermatheca, even if there was some sperm still in the uterus (Fig 4a-b). The exception to this trend was with intestinal expression of *lin-35(+)* in a *lin-35(n745)* mutant, which showed limited sperm localization to the spermatheca at 26°C. Overall, these data indicate that in *lin-35* mutant hermaphrodites raised at 20°C wildtype sperm can localize and should be able to fertilize oocytes. Therefore, there may be a defect in oocytes in *lin-35* mutants even at 20°C. However, at 26°C male sperm are able to localize to the spermatheca only if the *lin-35(+)* is expressed broadly in the soma, thus *lin-35* mutants have additional factors contributing loss of fertility at elevated temperatures.

**Figure 4:**
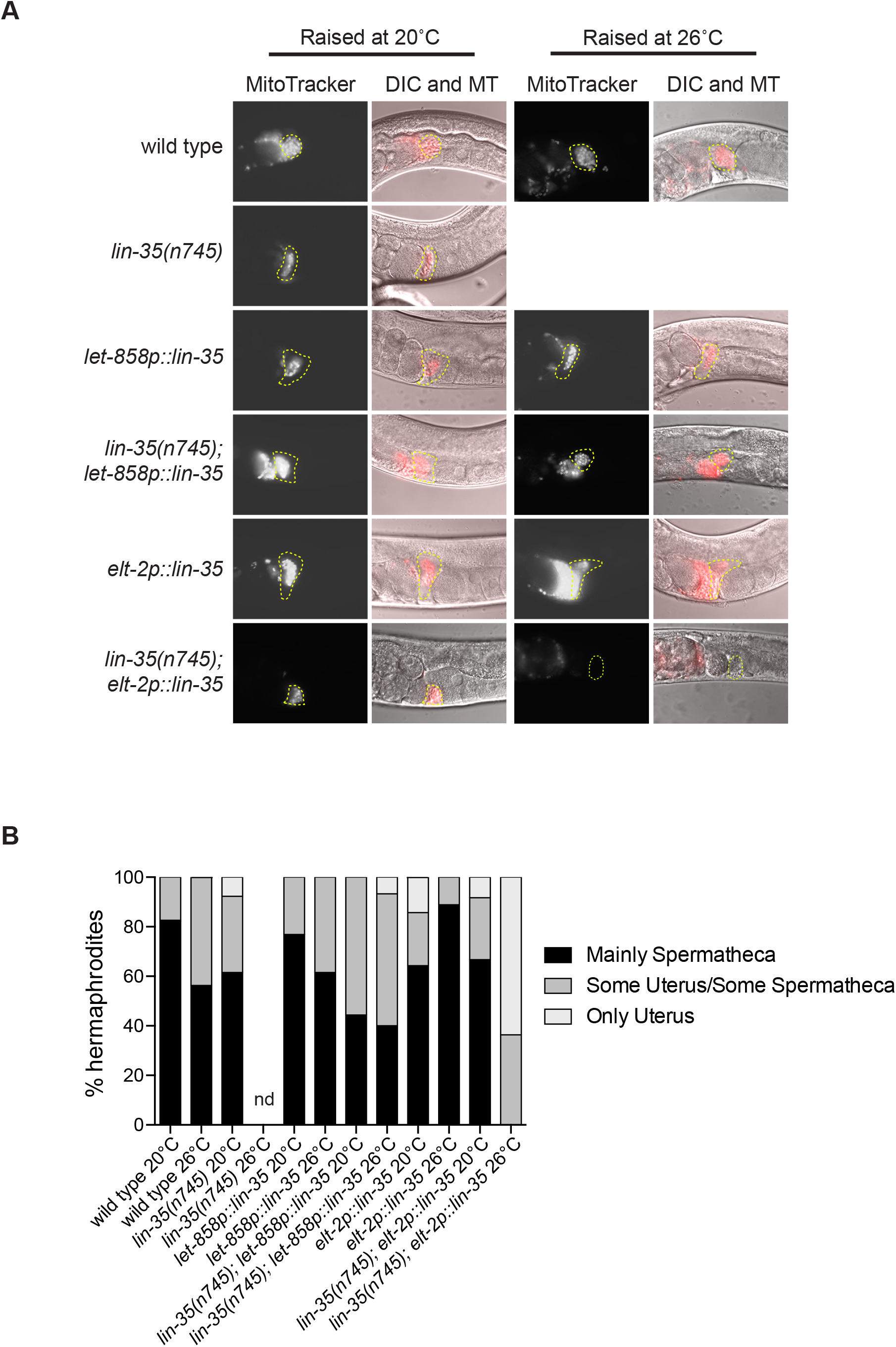
Male sperm do not migrate in *lin-35* mutants expressing the intestinal transgene at high temperature. (A) Representative images of spermatheca in hermaphrodites after mating with males stained with MitoTracker Red CMXRos. The location of spermatheca is indicated by yellow outlines. (B) Percentage of hermaphrodites that show different distributions of male sperm within the reproductive tract as either primarily within the spermatheca, a combination of sperm within the spermatheca and in the uterus, and sperm only within the uterus. Notably no *lin-35(n745); elt-2p::lin-35* hermaphrodites at 26°C have sperm in the spermatheca.

### Pan-somatic LIN-35(+) is necessary for oocyte formation at 26°C

To assess if the loss of fertility in *lin-35* mutants could be due to the inability of animals to produce oocytes, we assessed the number of oocytes in *lin-35(n745)* hermaphrodites with and without *lin-35(+)* transgenes at 20°C and 26°C. We scored each germline to determine if animals had more than one oocyte in a gonad arm (Fig 5a). We found that for all strains close to 100% of gonads had >1 oocyte with the exception *lin-35(n745)* with the intestinal *lin-35(+)* transgene at 26°C (Fig 5b). Germlines in *lin-35(n745); elt-2p::lin-35::GFP* hermaphrodites at 26°C generally looked small, had poor organization, and generally only had one cell proximal to the spermatheca that looked like an oocyte (Fig 5A). These data underscore the importance of broad somatic expression of *lin-35(+)* for maintaining germline function especially at 26°C.

**Figure 5:**
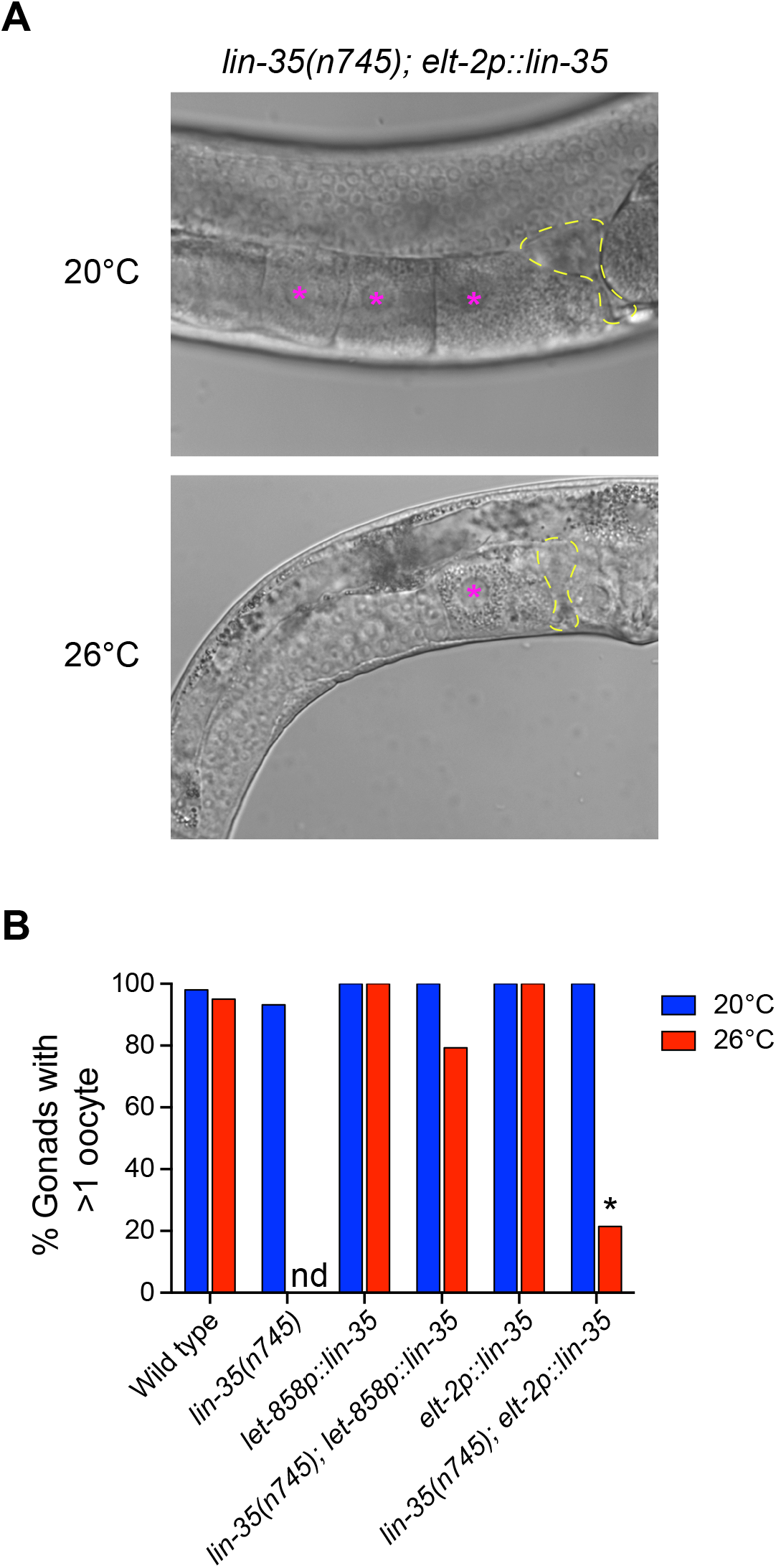
*lin-35* mutants expressing the intestinal transgene have fewer oocytes at high temperature. (A) Representative DIC images of *lin-35(n745); elt-2p::lin-35* hermaphrodite gonad at 20°C and 26°C. The location of spermatheca is indicated by yellow outlines and oocytes are represented by a pink asterisk. (B) The percentage of gonad arms in hermaphrodites that have >1 oocyte. Significantly fewer *lin-35(n745); elt-2p::lin-35* hermaphrodites at 26°C have >1 oocyte than at 20°C. * significantly different from same strain at 20°C. *P*-value ≤ 0.05. using Fisher’s exact test.

### PGL-1 localization to P granules is disrupted at 26°C in L1 larvae

Germline specific RNA-protein granules called P granules are known to be important for fertility, especially at 26°C [32]. At high temperature, certain protein components of P granules lose their localization to P granule condensates, including PGL-1 [33–34]. *lin-35* mutants display ectopic expression of germline genes, including PGL-1, in the intestine [7, 13]. We noticed that in some *lin-35* L1s at 26°C, there appeared to be mislocalization of PGL-1 within the two embryonic primordial germ cells (PGCs); therefore, we closely analyzed PGL-1 localization to P granules in L1 animals at both 20°C and 26°C. We found that for all strains at 20°C, the vast majority of L1 PGCs showed PGL-1 with a normal pattern of localization in puncta that are relatively evenly distributed around the nucleus of the two PGCs (Fig 6a-b). However, we found that for all strains, except *lin-35(n745); pie-1p::lin-35*, there was a significant increase in the number of L1s that had PGL-1 mislocalized within PGCs at 26°C compared to the same strain at 20°C (Fig 6b). The change in localization was never fully penetrant with some L1s showing completely normal PGL-1 localization (normal), some L1s showing a slight change in localization where there was a shift such that more PGL-1 localized the interface between the two PGCs (shifted central), and the most severe disruption where PGL-1 localized to generally only a few large puncta and there was often some diffuse PGL-1 staining (disorganized) (Fig 6a). While we saw no large differences between *lin-35(+)* and *lin-35* mutant strains, there was a strong trend for strains expressing extra LIN-35(+) solely in the PGCs to have fewer L1s display PGL-1 mislocalization. We also saw that in *lin-35* mutants, about 5% of larvae at elevated temperature only had one PGC (Fig 6a, bottom row). Therefore, while increased *lin-35* expression may be protective against disruption of P granules at elevated temperatures, disruption of P granules in L1 PGCs is a general temperature sensitive phenotype.

**Figure 6:**
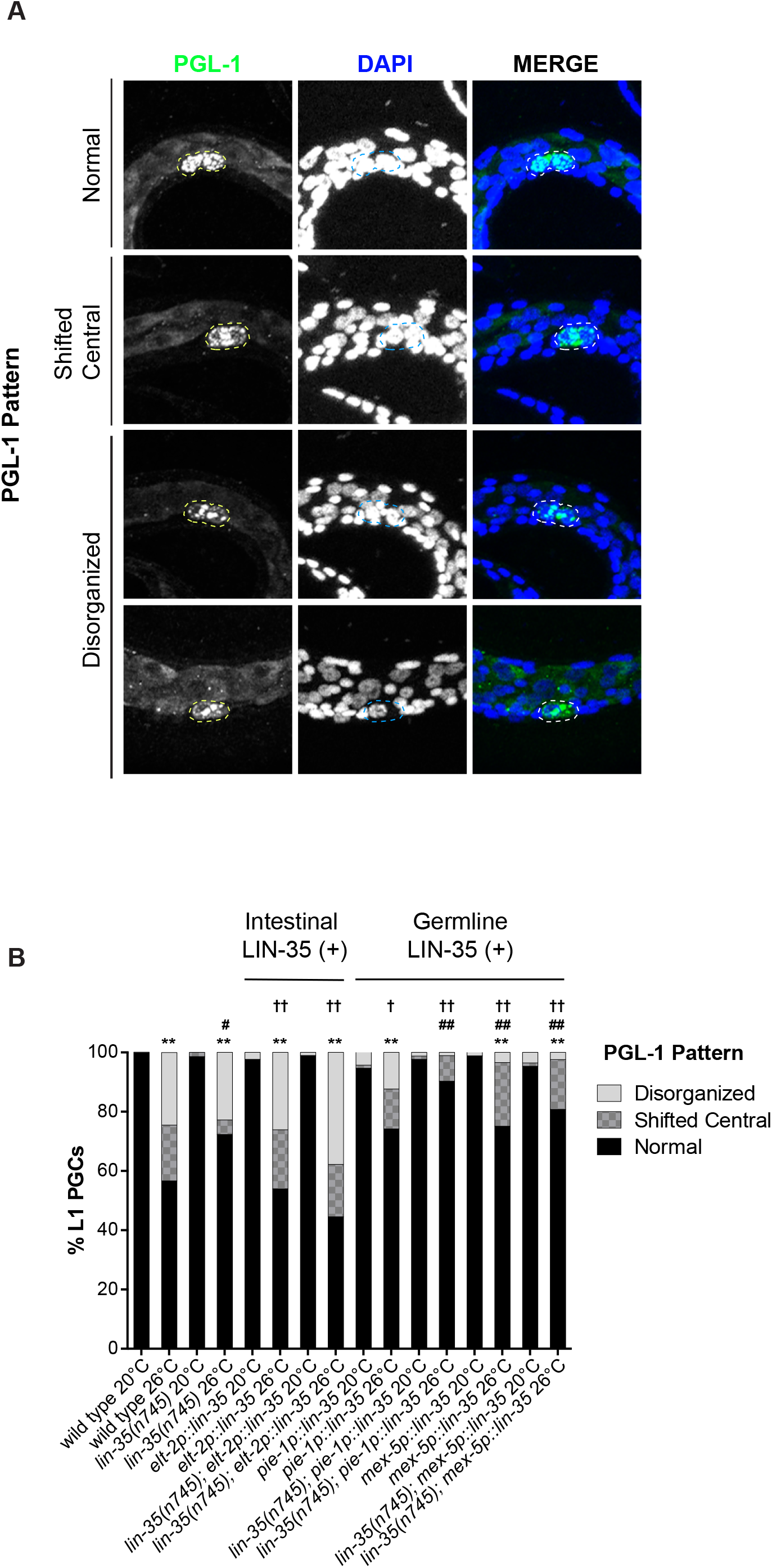
PGL-1 is mislocalized in L1 PGCs at high temperature. (A) Representative images showing patterns of PGL-1 localization in the two L1 primordial germ cells (PGCs) as “Normal” with evenly distributed puncta, “Shifted Central” with increased localization at the interface where the two PGCs meet, or “Disorganized” with only few, non-evenly distributed puncta/diffuse staining. All images are in *lin-35(n745)* at 26°C. (B) Distribution of PGL-1 localization patterns across different strains at different temperatures. Significantly more L1s showed PGL-1 staining not in the “Normal” pattern at 26°C compared to 20°C for most strains. ^*^ significantly different from same strain at 20°C, ^#^ significantly different from N2 at 26°C, ^†^ significantly different from *lin-35(n745)* at 26°C; ^†# *^ *P*-value < 0.05, ^††## **^ *P*-value ≤ 0.01 using Chi squared test using multivariable contingency table.

Previous work has also shown that PGL-1 can become diffuse in adult germlines subjected to short exposures of 29°C, specifically in the distal part of the germline near the mitotic zone and in oocytes [33]. We therefore looked to see if we could also see changes to PGL-1 localization in adult germlines of hermaphrodites that spent their entire life at 26°C. Unlike in L1 animals at 26°C, in wild type hermaphrodites we saw no significant diffuse or mislocalizated of PGL-1 in adult germlines (Table 1). In hermaphrodites expressing *lin-35(+)* in the intestine, *lin-35(n745); elt-2p::lin-35*, we also saw no significant changes in PGL-1 localization (Table 1). As expected, we saw oocyte defects, including endomitotic oocytes, in *lin-35(n745); elt-2p::lin-35* adult germlines at 26°C, but PGL-1 generally was found in punctate structures despite oocyte malformation. However, in hermaphrodites expressing *lin-35(+)* pan-somatically, *lin-35(n745); let-858p::lin-35*, we did see diffuse PGL-1 localization in the late pachytene and oocytes (Table 1). In adult germlines we never observed PGL-1 in larger puncta reminiscent of what we observed in the PGCs in L1 animals. This suggests that in the vast majority animals, the mislocalization of PGL-1 seen in L1 PGCs is resolved before adulthood and does not broadly contribute to the decrease in fertility seen at 26°C.

**Table 1:** Mislocalization of PGL-1 in Adult Germlines

| Strain | % Germlines with Diffuse PGL-1 (n) |  |  |
| --- | --- | --- | --- |
|  | Distal Region | Pachytene Region | Oocytes |
| Wild type 20°C | 0.0 (16) | 17.6 (17) | 0.0 (12) |
| Wild type 26°C | 0.0 (15) | 10.5 (19) | 0.0 (13) |
| *lin-35(n745)* 20°C | 19.2 (26) | 16.7 (36) | 4.3 (23) |
| *lin-35(n745);*<br>*let-858p::lin-35* 20°C | 0.0 (28) | 11.1 (18) | 0.0 (6) |
| *lin-35(n745);*<br>*let-858p::lin-35* 26°C | 5.6 (18) | 36.8 (19)* | 40.0 (15) |
| *lin-35(n745);*<br>*elt-2p::lin-35* 20°C | 0.0 (20) | 15.0 (20) | 7.7 (13) |
| *lin-35(n745);*<br>*elt-2p::lin-35* 26°C | 0.0 (10) | 9.1 (11) | 0.0 (10) |

\* significantly different from wild type at the same temperature, $P \leq 0.05$ , Fisher's exact test

## Discussion

Like other phenotypes in *lin-35* mutants, we show that fertility is temperature sensitive. We demonstrate that LIN-35 has a small role in the soma to protect fertility, but that germline expression LIN-35 is necessary for fertility, especially under moderate temperature stress. Interestingly, unlike most other LIN-35 functions, the role of LIN-35 in the germline appears to primarily be during post-embryonic development and dependent on zygotic and not maternal expression. Finally, LIN-35 in the soma contributes to proper oocyte formation and/or function under moderate temperature stress. There is a growing understanding that in addition to its other roles, LIN-35 plays an important role in a number of stress response pathways [35]. The work presented here underscores a new role in stress response for LIN-35 in regulating germline function at elevated temperature.

In our work we saw that somatic expression of LIN-35(+) weakly but significantly rescues fertility defects in *lin-35* mutants at 20°C. Progeny production relies on many somatic tissues including the intestine, somatic gonad, as well as neuronal input [14–15, 36]. *lin-35* mutants display a broad range of effects on different tissues, such as having generally have slow growth, so it is not surprising that somatic rescue of LIN-35(+) could have a positive effect on brood size. Wildtype function of LIN-35 in the intestine appears to be particularly important for fertility at 20°C given that the level of rescue is equivalent between solely intestinally expressed LIN-35(+) and pan-somatic expression of LIN-35(+). Previous research has shown that in *lin-35* mutant L1 animals, genes normally expressed in the intestine are disproportionately both up- and down-regulated [13]; therefore, the intestine appears to be one of the tissues most effected by loss of *lin-35*. However, when worms experience temperature stress at 26°C, there are marked differences in intestinally expressed LIN-35(+) and pan-somatic expression of LIN-35(+). This is easily seen with the smaller brood size and significantly fewer number of worms that are fertile in *lin-35* mutants expressing intestinal LIN-35(+) compared to worms expressing pan-somatic LIN-35(+) in either a wild type or *lin-35* mutant background. Additionally, at 26°C *lin-35* mutants expressing intestinal LIN-35(+) show significant defects in both sperm migration to the spermatheca and the number of oocytes present in the germline. While the somatic gonad is a clear support tissue necessary for fertility, expression of LIN-35(+) in both somatic gonad and intestine was not sufficient to rescue the low fertility seen in *lin-35* mutants. Which other somatic tissues require expression of LIN-35(+) to maintain fertility when animals are temperature stressed remain unclear. However, the reduced number of oocytes and the failure of sperm to migrate seen at high temperature may be related. Sperm migration to the spermatheca relies on polyunsaturated fatty acid (PUFA) signals that emanated from oocytes for proper targeting [37]. If oocytes are not forming or functioning properly this could directly lead to mislocalization of sperm and thus low fertility.

It is clear from our data that germline intrinsic expression of *lin-35(+)* is the primary source for promotion of fertility both under stress and non-stress temperature conditions. While there has been speculation about the various roles that LIN-35 may play in the germline, the role that has been clearly defined has been in promotion of germline apoptosis. *lin-35* mutants have a slightly lower level of physiological apoptosis under non-stress conditions [6], but importantly fail to increase the rate of germline apoptosis under various stress conditions, including DNA damage and starvation [6, 38]. Germline apoptosis has also been shown to be induced under specific elevated temperature conditions [19]. Therefore, a decreased level of germline apoptosis may contribute to the decreased fertility of *lin-35* mutants, especially under stress conditions. Our data suggests that *lin-35* mutant germline defects are due the loss of LIN-35 activity in either larval germline development or in the adult germline, as zygotic expression of LIN-35 can strongly rescue fertility at both low and high temperature. We also found that there is likely a defect in some aspect of oocyte formation or function as mating with wildtype males results in little to no rescue of fertility in *lin-35* mutants. Oocyte formation and/or function seems to be especially sensitive to temperature when LIN-35(+) is only present in the intestine.

Phenotypes of *lin-35* mutants are notably temperature sensitive. For example, *lin-35* mutant larvae only arrest 26°C, while ectopic expression of germline expressed genes in somatic tissues and the synthetic multivulva phenotype seen in *lin-35* mutants are enhanced with increased temperature [13,20]. Here we show that both brood size and oocyte formation in *lin-35* mutants is temperature sensitive, expanding the temperature sensitive nature of *lin-35* phenotypes to the germline. Historically temperature sensitive phenotypes have been most clearly explained in situations where a hypomorphic allele of a gene results in a protein that becomes destabilized at increased temperatures. However, the *lin-35(n745)* allele contains premature stop codon, has no visible protein expression, and has generally been categorized as a putative null allele [1]. Therefore, it seems unlikely that loss of functional LIN-35 protein is what ties these temperature sensitive phenotypes together. Another option could be that any protein complex that LIN-35 normally interacts with becomes unstable at high temperature when LIN-35 is absent. The most well characterized complexes that LIN-35 is a part of are the E2F complex and the DREAM complex [39–40]. Previous work has shown that loss of other members of the E2F complex has distinctly different germline phenotypes and leads to different changes in gene expression in mutant germlines [12]. There have been no studies to date that have investigated the formation of the DREAM complex in the germline, but in the soma there is a significant but not complete loss of DREAM complex binding to its targets in *lin-35* mutants even at 20°C [27]. There could be increased loss of the DREAM complex binding in the soma at 26°C, but this has yet to be explored. It has been shown that there are increased fertility defects in double mutants between members of the MuvB core of the DREAM complex and *lin-35* [27]. We have also shown that loss of *lin-35* or the MuvB core member *lin-54* in embryos leads to temperature sensitive changes in compact chromatin formation that likely underlie the temperature sensitivity of ectopic germline gene expression and the high temperature larval arrest phenotypes [21]. Together these data indicate that there is likely an interaction of LIN-35 with at least the MuvB core of the DREAM complex in germline that is important for germline function. Additionally, we speculate that interaction between the MuvB core and its targets or the function of the complex is interrupted by increased temperature in *lin-35* mutants.

---

**Figure S1:**
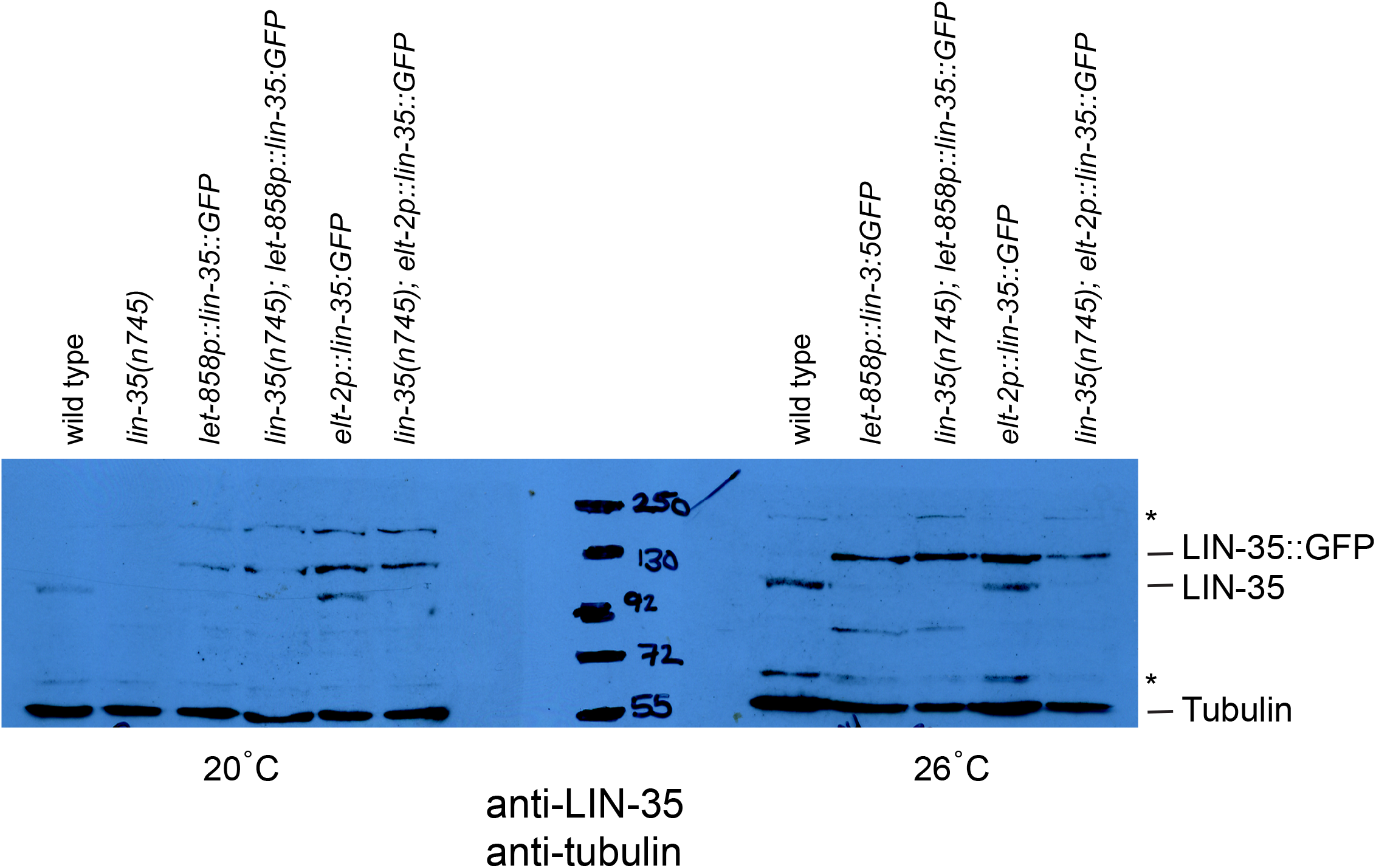
*let-858* driven expression of LIN-35::GFP represses expression of wildtype LIN-35 protein. Western blot using anti-LIN-35 antibodies shows expression of both the wildtype LIN-35 protein and LIN-35::GFP tagged protein expressed from somatic transgenes from worms grown at either 20°C or 26°C. In wildtype worms containing the *let-858p::lin-35::GFP* transgene, only LIN-35::GFP protein can be seen, while in wildtype worms containing the *elt-2p::lin-35::GFP* transgene, both LIN-35 and LIN-35::GFP protein can be seen. anti-Tubulin was used on the same blot as a loading control. * non-specific background bands.

## REFERENCES

1. Lu X, Horvitz HR. *lin-35* and *lin-53*, two genes that antagonize a *C. elegans* Ras pathway, encode proteins similar to Rb and its binding protein RbAp48. Cell. 1998 Dec 23; 95(7):981–91.

2. Classon M, Dyson N. p107 and p130: versatile proteins with interesting pockets. Exp. Cell Res.2001 Mar 10;264(1):135–147.

3. Julian LM, Blais A. Transcriptional control of stem cell fate by E2Fs and pocket proteins. Front Genet. 2015 Apr 28; 6:161.

4. Grishok A, Sharp PA. Negative regulation of nuclear divisions in *Caenorhabditis elegans* by retinoblastoma and RNA interference-related genes. PNAS. 2005 Nov 29;102(48):17360–17365.

5. Ouellet J, Roy R. The *lin-35/Rb* and RNAi pathways cooperate to regulate a key cell cycle transition in *C. elegans*. BMC Dev Biol. 2007 Apr;7: 38.

6. Schertel C, Conradt B. *C. elegans* orthologs of components of the RB tumor suppressor complex have distinct pro-apoptotic functions. Development. 2007 Oct;134(20):3691–701.

7. Wang D, Kennedy S, Conte Jr D, Kim JK, Gabel HW, Kamath RS, et al. Somatic misexpression of germline P granules and enhanced RNA interference in retinoblastoma pathway mutants. Nature. 2005 Jul 28;436(7050):593–7.

8. Bender AM, Wells O, Fay DS. *lin-35/Rb* and *xnp-1/ATR-X* function redundantly to control somatic gonad development in *C. elegans*. Dev. Biol. 2004 Sept 15; 273(2): 335–349.

9. Ferguson EL, Horvitz HR. The multivulva phenotype of certain *Caenorhabditis elegans* mutants results from defects in two functionally redundant pathways. Genetics. 1989 Sep; 123(1):109–121.

10. Reddien PW, Andersen EC, Huang MC, Horvitz HR. DPL-1 DP, LIN-35 Rb and EFL-1 E2F act with MCD-1 zinc-finger protein to promote programmed cell death in *Caenorhabditis elegans*. Genetics. 2007 Apr;175(4):1719–1733.

11. Fay DS, Keenan S, Han M. *frz-1* and *lin-35/Rb* function redundantly to control cell proliferation in *C. elegans* as revealed by a nonbiased synthetic screen. Genes Dev. 2002 Feb 15;16(4):503–517.

12. Chi W, Reinke, V. Promotion of oogenesis and embryogenesis in the *C. elegans* gonad by EFL-1/DPL-1(E2F) does not require LIN-35(pRB). Development. 2009 May-Jun; 133(5-6):3147–3157.

13. Petrella LN, Wang W, Spike CA, Rechtsteiner A, Reinke V, Strome S. synMuv B proteins antagonize germline fate in the intestine and ensure *C. elegans* survival. Development. 2011 Mar;138(6):1069–1079.

14. Geisler F, Coch RA, Richardson C, Goldberg M, Bevilacqua C, Prevedel R, et al. Intestinal intermediate filament polypeptides in *C. elegans:* Common and isotype-specific contributions to intestinal ultrastructure and function. Sci Rep. 2020 Feb 21;10(1):3142.

15. Hubbard EJ, Greenstein D. The *Caenorhabditis elegans* gonad: a test tube for cell and developmental biology. Dev Dyn. 2000 May;218(1):2–22.

16. Begasse ML, Leaver M, Vazquez F, Grill SW, Hyman AA. Temperature dependence of cell division timing accounts for a shift in thermal limits of *C. elegans* and *C. briggsae*. Cell Rep. 2015 Feb 10; 10(5):647–653.

17. Harvey SC, Viney ME. Thermal variation reveals natural variation between isolates of *Caenorhabditis elegans*. J Exp Zool B Mol Dev Evol. 2007 July 15;308(4):409–416.

18. Petrella LN. Natural variants of *C. elegans* demonstrate defects in both sperm function and oogenesis at elevated temperatures. PLoS One. 2014 Nov 7;9(11):e112377.

19. Poullet N, Vielle A, Gimond G, Ferrari C, Braendle C. Evolutionarily divergent thermal sensitivity of germline development and fertility in hermaphroditic *Caenorhabditis* nematodes. Evol Dev. 2015 Nov-Dec;17(6):380–97.

20. Andersen EC, Saffer AM, Horvitz HR. Multiple levels of redundant processes inhibit *Caenorhabditis elegans* vulval cell fates. Genetics. 2008 Aug;179(4):2001–2012.

21. Costello ME, Petrella LN. *C. elegans* synMuv B proteins regulate spatial and temporal chromatin compaction during development. Development. 2019 Oct 9;146 (19):dev174383.

22. Latorre I, Chesney MA, Garrigues JM, Stempor P, Appert A, Francesconi M, et al. The DREAM complex promotes gene body H2A.Z for target repression. Genes Dev. 2015 Mar 1; 29(5):495–500.

23. Strome S, Wood WB. Generation of asymmetry and segregation of germ-line granules in early C. elegans embryos. Cell. 1983 Nov; 35(1):15–25.

24. Kawasaki I, Shim YH, Kirchner J, Kaminker J, Wood WB, Strome S. PGL-1, a predicted RNA-binding component of germ granules, is essential for fertility in C. elegans. Cell. 1998 Sep 4;94(5):635–45.

25. Kelly WG, Fire A. Chromatin silencing and the maintenance of functional germline in *Caenorhabditis elegans*. Development. 1998 Jul;125(13):2451–6.

26. Rios C, Warren D, Olson B, Abbott AL. Functional analysis of microRNA pathway genes in the somatic gonad and germ cells during ovulation in *C. elegans*. Dev Biol. 2017 Jun 1;426(1)115–125.

27. Goetsch PD, Garrigues JM, Strome S. Loss of the *Caenorhabditis elegans* pocket protein LIN-35 reveals MuvB’s innate function as the repressor of DREAM target genes. PloS Genet. 2017 Nov 1;13(11) e1007088.

28. Kudron M, Niu W, Lu Z, Wang G, Gerstein M, Snyder M, et al. Tissue-specific direct targets of *Caenorhabditis elegans* Rb/E2F dictate distinct somatic and germline programs. Genome Biol. 2013 Jan 23;14(1):R5.

29. Edgley ML, Baillie DL, Riddle DL, Rose AM. Genetic balancers. Wormbook. 2006 Apr 6:1–32.

30. Ward S, Carrel JS. Fertilization and sperm competition in the nematode *Caenorhabditis elegans*. Dev Biol. 1979 Dec;73(2)304–21.

31. Han SM, Cottee PA, Miller MA. Sperm and oocyte communication mechanisms controlling *C. elegans* fertility. Dev Dyn. 2010 May;239(5):1265–81.

32. Voronina E. The diverse functions of germline P-granules in *Caenorhabditis elegans*. Mol Reprod Dev. 2013 Aug; 80(8):624–31.

33. Fritsch AW, Diaz-Delgadillo AF, Adame-Arana O, Hoege C, Mittasch M, Kreysing M, et al. Local thermodynamics govern formation and dissolution of *Caenorhabditis elegans* P granule condensates. Proc Natl Acad Sci USA. 2021 Sept 14;118(37):e2102772118.

34. Putnam A, Cassani M, Smith J, Seydoux G. A gel phase promotes condensation of liquid P granules in *Caenorhabditis elegans* embryos. Nat Struct Mol Biol. 2019 Mar; 26(3):220–226.

35. Gonzalez-Rangel AA, Navarro RE. LIN-35 beyond its classical roles: its function in the stress response. Int J. Dev Biol. 2021;65(4-5-6):377–382.

36. Cottee PA, Cole T, Schultz J, Hoang HD, Vibbert J, Han SM et al. The *C. elegans* VAPB homolog VPR-1 is a permissive signal for gonad development. Development. 2017 Jun 15; 144(12):2187–2199.

37. Ellis RE, Stanfield GM. The regulation of spermatogenesis and sperm function in nematodes. Semin Cell Dev Biol. 2014 May;29:17–30.

38. Lascarez-Legunas LI, Silva-Garcia CG, Dinkova TD, Navarro RE. LIN-35/Rb causes starvation-induced germ cell apoptosis via CED-9/Bcl2 downregulation in *Caenorhabditis elegans*. Mol. Cell Biol. 2014 Jul;34(13):2499–2516.

39. Ceol CJ, Horvitz HR. *dpl-1* DP and *efl-1* E2F act with *lin-35* Rb to antagonize Ras signaling in *C. elegans* vulval development. Mol. Cell 2001 Mar;7(3):461–473.

40. Harrison MM, Ceol CJ, Lu X, Horvitz HR. Some *C. elegans* class B synthetic multivulva proteins encode a conserved LIN-35 Rb-containing complex distinct from a NuRD-like complex. Proc Natl Acad Sci USA. 2006 Nov 7;103(45):16782–7.

